# Enhancing sensitivity and controlling false discovery rate in somatic indel discovery using a latent variable model

**DOI:** 10.1101/121954

**Authors:** Louis J. Dijkstra, Johannes Köster, Tobias Marschall, Alexander Schönhuth

**Affiliations:** Universiteit van Amsterdam, Amsterdam, The Netherlands; Centrum Wiskunde & Informatica, Amsterdam, Netherlands; Dana-Farber Cancer Institute, Harvard Medical School, Boston, Massachussetts; Max Planck Institute for Informatics, Saarbrücken, Germany; University Saarbrücken, Germany

## Abstract

Cancer is a genetic disorder in the first place. Therefore, next-generation sequencing (NGS) based discovery of somatically acquired genetic variants has gained widespread attention. Computational prediction of somatic variants, however, is affected by a variety of confounding factors. In addition to the uncertainties that one commonly encounters also in germline variation prediction, such as misplaced and/or inaccurate read alignments, cancer heterogeneity and impure samples significantly add to the issues. Overall, this hampers state-of-the-art indel discovery tools to discover somatic indels at operable performance rates, although they perform excellently when calling germline indels. While affecting all size ranges, both common and cancer-specific problems interfere in particularly unfavorable ways in the prediction of somatic midsize (30-150 bp) insertions and deletions.

Here, we present a latent variable model that can take the major confounding factors and uncertainties into a unifying account. Using this modeling framework, we *first* demonstrate how to *efficiently* compute the probability for a (putative) indel to be somatic, thereby resolving a principled computational runtime bottleneck in Bayesian uncertainty quantification. *Second*, we show how to reliably estimate the allele frequencies for a given list of indels. *Third*, we also present an intuitive and effective way to control the false discovery rate, an issue in genetic variant discovery that has been found notoriously hard to deal with. As a tool that implements all methodology developed, we present PROSIC (PROcessing Somatic Indel Calls). PROSIC achieves significant improvements in particular in terms of recall when applied to deletion call sheets, as provided by prevalent state-of-the-art tools, in comparison to their integrated somatic indel calling routines.

The software is publicly available at https://prosic.github.io and can be easily installed via https://bioconda.github.io.

## Introduction

Cancer is a genetic disorder in the first place; somatic mutations turn originally healthy cells into a heterogeneous mix of aberrantly evolving cell clones [2]. Empowered by the routine application of newer generation sequencing technologies [7], global consortia [26] have launched petabyte scale projects concerned with the discovery and annotation of somatic mutations in cancer genomes [28]. Benefits of such systematic analysis of somatic mutations include improved diagnosis, staging and therapy protocol selection in the clinic.

Still, the most prevalent approach to somatic mutation discovery are bulk re-sequencing protocols. Next-generation sequencing (NGS) fragments of a cancer genome and a matched healthy (a.k.a. *control*) genome are aligned against the reference genome. Somatic mutations are discovered by comparing variants detected in the two genomes; those present in the cancer genome, but not in the control genome, are output as somatic.

This at first glance simple looking differential analysis is complicated by several factors, which add to the usual issues arising in non-differential settings. *First*, cancer heterogeneity plays a particularly disturbing role. While in germline variant discovery, variants come at allele frequencies of either 0.0, 0.5 or 1.0, reflecting absence, heterozygosity or homozygosity respectively, the usually unknown clonal structure allows no prior assumptions in somatic variant discovery. Since each of the clones is characterized by their own set of somatic variants, some of which are shared with other clones and some of which are unique, any allele frequency between 0 and 1 may apply for the bulk of somatically varied cells. *Second*, the tumor sample usually contains a non-negligible amount of healthy cells. Estimating the *level of purity* or *cellularity*, i.e., the fraction of cancer cells present in the sample, remains difficult, although recent progress has been made [8]. It is therefore advantageous to take purity into account.

Related complications become particularly disturbing when aiming at the discovery of insertions and deletions of of 30 - 120 bp in length^1^. This size range has been termed *NGS twilight zone of indels* [17, 27]. The reason is that alignment data associated with such indels are particularly uncertain. Thereby, two major classes of uncertainties apply.

(1) *Alignment uncertainty:* Alignments of fragments affected by indels longer than 30 bp may easily be misplaced by short read alignment tools. In particular, the combination of indels and repeat elements can interfere in unfavorable ways [29]. This adds to traditional issues when dealing with potentially gapped sequence alignments (see e.g. [16] for prominent artifacts such as gap wander, gap annihilation, and so on).
(2) *Typing uncertainty*: Even if placed correctly, the alignments may not give rise to clearly discernible variant signals. For example, fragment length considerations become statistically more involved for midsize indels [17]. In comparison to short indels, uncertain gap placement within the alignments (see above) plays a significant role.

So, “twilight zone” indel calling requires particular precautions with respect to uncertain data handling, as has been noted in various places [12, 17, 18, 27]. There are good tools for the discovery of somatic single nucleotide variants (see e.g. [1, 9, 11, 25]). Tools presented for discovery of somatic insertions and deletions, such as [23, DELLY], [24, PLATYPUS], [30, PINDEL] and (most recently) [22, LANCET] are very conservative, one reason being that they ignore the majority of uncertain data signals, which results in significantly reduced recall rates.

*Here, we present a Bayesian latent variable model that takes the major disturbing data uncertainties into account*. As is usual in uncertain data analysis, the major computational bottleneck of the analysis is the exponential amount of possibly correct data scenarios. For example, when evaluating *n* alignments that provide information about a putative somatic variant at a particular locus, we must take into account that each of the *n* alignments could be (a) misplaced, or, if placed correctly could either (b) be affected by the variant or (c) not be affected—only one of these options is correct. This induces 3*^n^* possible scenarios of correct interpretation of the alignment data; too many, because *n*, the fragment coverage of the locus, typically ranges between 30 and 50. Here, we provide discovery algorithms that run in time linear—and not exponential—in the coverage, a major methodological contribution of this work.

As for the application of our model in somatic variant discovery, *the idea of this work is to turn generic indel callers into high-performance somatic indel discovery engines on the one hand and to significantly increase sensitivity for stand-alone somatic indel discovery tools*. At this point, we do not aim to devise stand-alone indel discovery tool. Our idea is to provide a sound statistical framework for enhancing related performance rates. As one example of the first class consider PINDEL [30] which has turned over the years into a highly engineered, non-differential indel discovery tool. However, when using this high-performance discovery machine for somatic indel discovery, performance rates substantially drop. As an example for the latter class consider DELLY [23], since recently also a high-performance somatic indel discovery tool. As we will demonstrate, postprocessing somatic indel calls from DELLY with PROSIC leads to a relative increase of more than 60-70% in recall across all indel size ranges, without loosing precision. We emphasize another time that this is a statistically involved undertaking. Here, we resolve the inherent issues. To the best of our knowledge, related statistical machinery—which, as we feel, are of great practical value—has not yet been presented in the literature.

*As our major result*, we are indeed able to significantly increase recall rates when discovering somatic deletions, while being on a par with, if not improving on the precision achieved when making use of ad-hoc routines for somatic variant discovery implemented by the callers themselves, which sometimes, thanks to highly engineered (while still ad-hoc) filtering procedures, are truly excellent. Thereby, improvements show in particular for somatic twilight zone deletions (here: 30-250bp). Last but not least, our framework gives rise to a natural and intuitive procedure that allows to control the false discovery rate (FDR) in somatic indel discovery experiments, a notoriously difficult issue in genetic variant discovery in general (see [3, 17] for controlling FDR in germline indel discovery).

## Methods

### 2.1 Notation, Approach and Objectives

We denote *observable variables* by (Latin) capital letters (e.g. *Z*). *Realizations* of these variables are denoted by small (Latin) letters (e.g. *z*). *Hidden/latent variables* are denoted by (small) Greek symbols. Vectors are denoted by boldface letters (e.g. ***Z*** = (*Z*_1;_*Z_k_*) or *z* = (*z*_1,…,_*z_k_*). We use super-/subscripts *h* and *t* for the healthy and the tumor sample and *c* to only refer to cancer cells. Note that the tumor sample also contains healthy cells, which is usually referred to as *impurity;* let 0 ≤ *α* < 1 be the fraction of healthy cells in the tumor sample. Let us fix a particular variant locus; we then denote the relevant alignment data (encoding alignment length and/or gap content, see subsection 2.2 for details) in the healthy and the tumor sample by 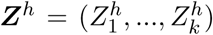 and 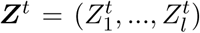, where each of the 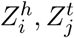, *i* = 1,*k, j* = 1,…, *l* represents one alignment covering the fixed variant locus. By *variant allele frequency* (*VAF*), we refer to the fraction of genome copies in the sample affected by the variant. We denote this (unknown) frequency in the healthy and the tumor sample by *θ_h_* and *θ_t_*, respectively. Since healthy cells are diploid, we can restrict *θ_h_* to the values 0, ^1^/2 and 1 corresponding to absence, heterozygosity and homozygosity. Prior such knowledge about *θ_t_* is usually unknown at the time of analysis, as it would require to understand the clonal structure among the cancer cells. So we let *θ_t_* vary over the entire unit interval [0, 1]. For *θ_c_*, defined to be the VAF among only the cancer cells, we obtain the relationship 
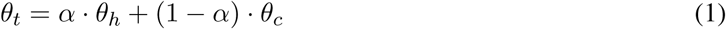
which also varies over the entire unit interval [0,1].

We fix a (putative) variant locus and consider the alignment data ***Z**^h^, **Z**^t^* of alignments covering the locus in the healthy and the tumor sample. Our objective is to *efficiently compute* 
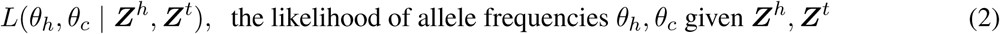

To understand the difficulties, consider that each of the alignments, 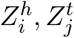 could either be (a) incorrect, (b) correct and not affected by the variant or (c) correct and affected by the variant. We recall that there is considerable uncertainty about this for alignments at *midsize* indel loci in particular. Following a *fully Bayesian approach to inverse uncertainty quantification* [15], we attach hyperparameters 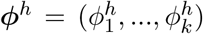, 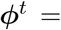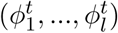 to the alignments ***Z**^h^, **Z**^t^* where each of the 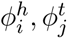 is ternary-valued, taking values in {0,1,2}, reflecting the above-mentioned three cases (a), (b) and (c). For example, 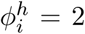 reflects that 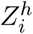 is correct and associated with the variant, because, for instance, alignment 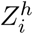 has a high alignment score and exhibits a gap agreeing with the coordinates of the putative indel under consideration. Computation of 
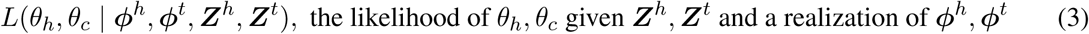
is straightforward, because once realizations of *ϕ^h^, ϕ^t^* have been specified, ***Z**^h^, **Z**^t^* are no longer uncertain. Note further that probability distributions *P*(*ϕ^h^, ϕ^t^*) can be obtained from the alignment tool. See subsection 2.2 below for details on these points.

Encouraged by these observations, we straightforwardly compute 
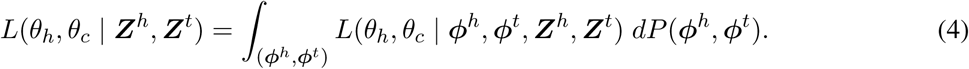

However, there are 3*^k^*^+^*^l^* different choices for realizations of *ϕ^h^, ϕ^t^*. So, computation of the integral requires *O*(3*^k^*^+^*^l^*) runtime, meaning that it is exponential in the fragment coverage of the variant locus.

While fully Bayesian approaches to inverse uncertainty quantification are certainly desirable, the integration over uncertainty hyperparameters constitutes their principled computational bottleneck. Here, we can overcome this bottleneck; the following is a *main result of this treatment*.

#### Result

The integral (4) can be evaluated in *O*(*k* + *l*) runtime, with a small constant factor.

This insight renders computing (2) and (4) tractable for all putative indel loci in a cancer genome. The result follows from Theorem 2.2, see Corollary 1 at the end of subsection 2.2.

##### Application I: Classification

Exploiting the efficiency in computing (2), we can determine posterior probabilities for a given indel to be either somatic, a germline variant or absent. The posterior probabilities of each of these cases can be translated into a statement about the values of *θ_h_* and *θ_c_*, see Figure 1a. In the case of a somatic indel, the variant does not occur among the healthy cells, i.e. *θ_h_* = 0, while it is present among the cancer cells, i.e. *θ_c_* ∈ (0,1]. Germline variants are present in the healthy cells, i.e. *θ_h_* ∈ {^1^*/*2,1} while the particular choice of *θ_c_* is irrelevant, i.e. *θ_c_* ∈ [0, 1]. Finally, if the indel is absent, both VAFs are zero: *θ_h_* = 0 and *θ_c_* = 0. We can compute the posterior probabilities for these three cases as [see also Subsections 2.2 and Appendix A for further details] 
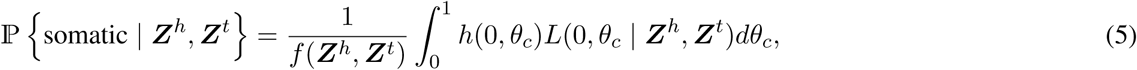

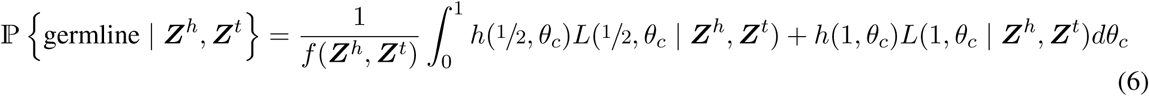
and 
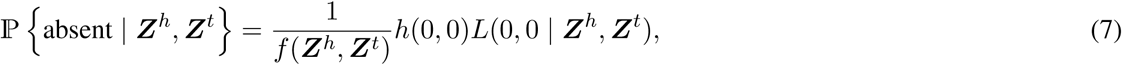
where L(·, · | ***Z**^h^, **Z**^t^*) is the likelihood function from eq. (2), *h*(*θ_h_, θ_c_*) is a prior probability for the given combination of allele frequencies, and 
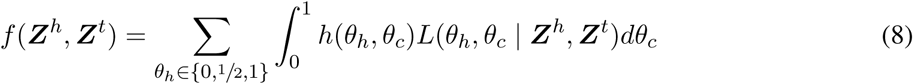
is the marginal probability of the data. Here, we assume no further information on the clonal structure of the tumor sample, and use a uniform prior, such that the *h*(*θ_h_, θ_c_*) cancels out from above equations. Note that the prior *h*(*θ_h_, θ_c_*) allows to integrate prior knowledge about zygosity rates (for germline variants) and clonal structure (for somatic variants) if available in the future. The integrals are numerically approximated using the quadrature rule. Note that *key to success of numerical approximation of these integrals is the efficient computation* of L(·, · | ***Z**^h^, **Z**^c^*), as warranted by Corollary 1 below. This approach yields a posterior probability for all three categories. These probabilities can be used for filtering output, e.g. fix a threshold *τ* and output all indels as somatic where (5) is greater than *τ*.

**Figure 1:**
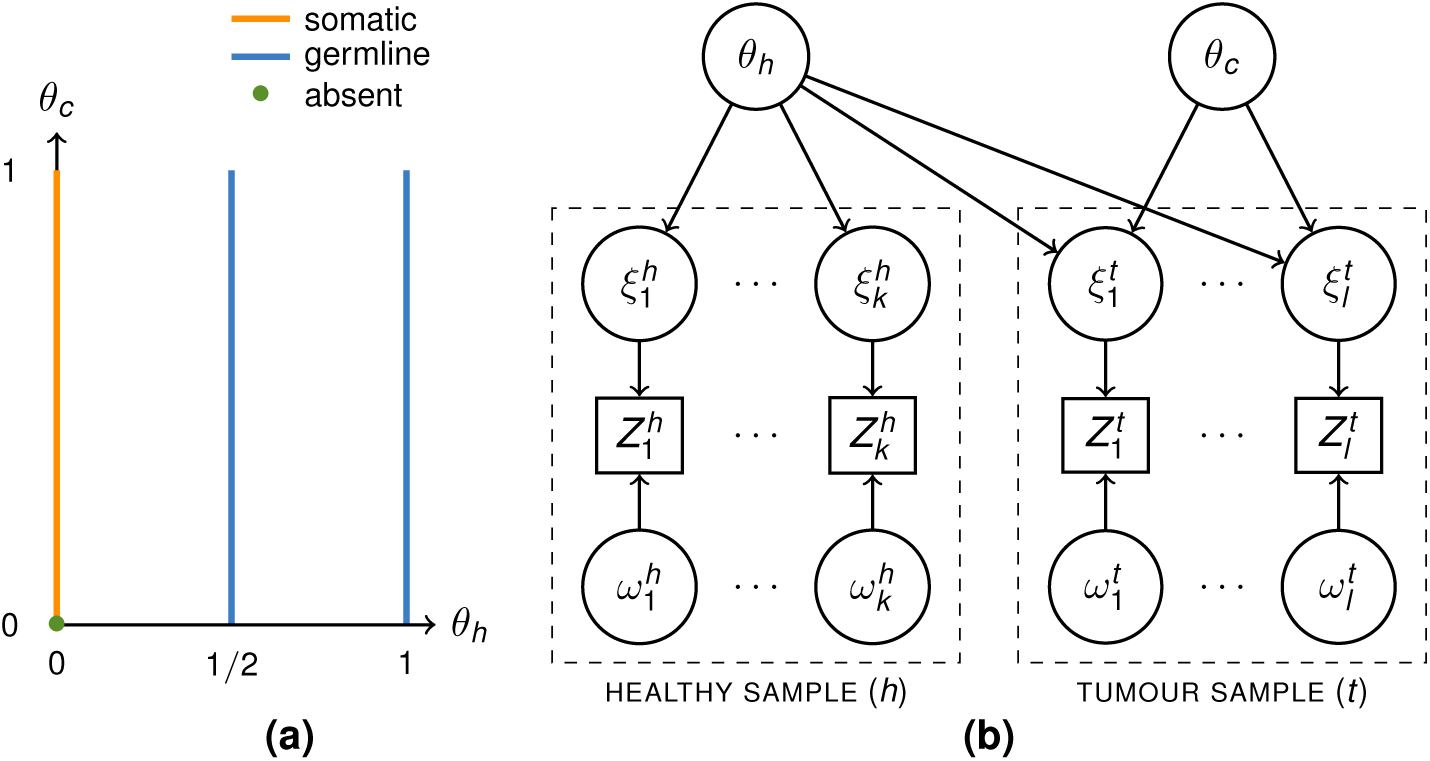
(**a**) A visualization of the parameter space Θ of the VAFs. *Orange:* somatic variants agree with (*θ_h_, θ_c_*) ∈ {0} × (0,1], which means that no healthy cells have the variant (*θ_h_* = 0), while some cancer clones do have the variant (*θ_c_* > 0). By analogous considerations we find germline variants (*blue*) described by 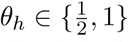 and absent variants (*green dot*) by *θ_h_* = 0, *θ_C_* = 0. (**b**) A diagram of the model presented in Section 2.2 with all its variables (circles=latent; rectangles=observable). Each column corresponds to one alignment 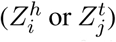 with its hyperparameters 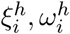 or 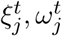. Due to (potential) sample impurity (denoted by *α* in the text), *θ_h_* has an influence on the alignments 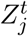 from the tumor sample.

##### Application II: Variant allele frequency estimation

The general model presented in section 2.2 allows us to estimate the VAFs of the healthy and cancer cells: the maximum a posteriori (MAP) estimate of *θ_h_* and *θ_c_* is that point in the parameter space Θ (see Figure 1a) for which the posterior distribution 
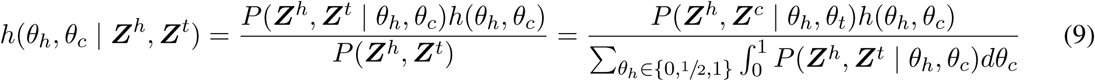
attains its maximum. We again assume a uniform prior *h*(*θ_h_, θ_c_*) over the parameter space Θ (see Figure 1a). Hence, *h*(*θ_h_, θ_c_* | ***Z**^h^*, ***Z**^t^*) agrees with the likelihood function and the maximum 
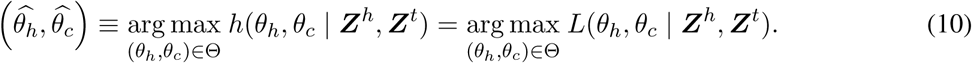
is the maximum likelihood estimate (MLE). The likelihood function in eq. (2) is a higher-order polynomial in *θ_h_* and *θ_c_* as follows from the computations in Appendix A which makes it infeasible to derive its maximum analytically. We can nevertheless prove the following helpful theorem.

### Theorem 2.1

*For fixed θ*_h_* the logarithm of the likelihood function θ*_c_ → L*(θ*_h_*, θ*_c_ | Z^h^, Z^t^*) is concave on the unit interval 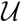 = [*0, 1*]. Hence the likelihood function attains a unique global maximum 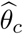 on* [0, 1].

See Theorem B.1 in Appendix B for a detailed technical exposition, including a proof and a list of extra conditions for this theorem necessary to hold, all of which apply in practice. Since the loglikelihood function is strictly concave, the required maxima can be easily determined numerically.

### Application III: False Discovery Rate (FDR) Control

After annotating all putative indels with a probability (5) to be somatic, it remains to filter this list to create reasonable output. Thereby, a common goal is control the false discovery rate, the expected relative amount of mistaken predictions. FDR control in variant discovery is of importance for various reasons. However, it has been found notoriously difficult to deal with in the literature so far (see [3, 17] for examples of FDR control procedures in variant discovery); the vast majority of discovery tools do not allow for such control. Our framework offers an intuitive and effective remedy.

Namely, we swap the roles of control and cancer genome in all steps of the discovery procedure. This yields a list of indels (*control indels*), all of which either reflect germline indels or artifacts, annotated with probabilities (5) to be somatic (*control probabilities*). Because none of the indels is a true discovery, sampling control probabilities reflects to sample probabilities to be somatic from the null hypothesis, that is for germline indels or alignment artifacts.

Returning to the original indels and their probabilities (5) to be somatic, we can compute p-values for each indel. We determine this p-value as the probability that a randomly sampled value from the null hypothesis distribution is at least as large as the probability to be somatic for the indel in question. Upon computation of a p-value for each original indel, we can sort these p-values and apply the Benjamini-Hochberg procedure to control for a given FDR *β*.

In a final remark, we found Bayesian type FDR control (note that our setting is perfectly Bayesian), as outlined for example in [21], to not work sufficiently well. A likely reason is that the null hypothesis distribution violates some consistency assumptions necessary to hold, which can be attributed to systematic alignment artifacts.

### 2.2 The Model

We present a graphical model that captures all dependency relationships among the variables relating to the computation of *L*(*θ_h_, θ_c_* | ***Z***^h^, ***Z**^t^*) while taking all major uncertainties into account. See Figure 1(**b**) for this graphical model.

#### Hyperparameters, Dependencies and Distributions

Beyond observable variables for alignment data 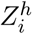, *i* = 1,…, *k*, 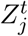 *j* = 1,…, *l* and latent variables *θ_h_, θ_c_* for allele frequencies, we introduce latent variables,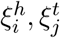 ∈ {0, 1} to specify whether alignments 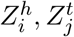 are associated with the variant 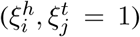 or not 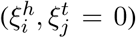 and 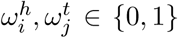 to model correct placement of alignments 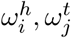 = 1 if correct and 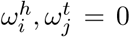, otherwise). Together, the latent variables 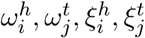, specify the ternary valued hyper-parameters 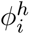, 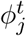 from subsection 2.1 above, where *ϕ* = 0, *ϕ* = 1 and *ϕ* = 2 correspond to {*ω* = 0}, {*ω* = 1,ξ = 0} and {*ω* = 1,ξ = 1}, respectively. For example 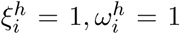 translates into 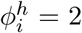, the case that alignment 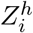 is correct and associated with the variant. Note that the case *ω* = 0, which indicates incorrectness of the alignment, renders specification of other variables obsolete, because this particular alignment cannot provide information about the variant. Of course, knowledge about realizations of these hyperparameters is unknown at the time of the analysis, so these variables remain hidden.

As just outlined, we model the two major sources of uncertainty, 1) *alignment uncertainty* and 2) *typing uncertainty*, by associating every observation 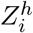, 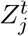 with two binary-valued, latent variables, reflecting uncertainty hyperparameters, 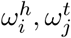 and 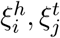 *First*, 
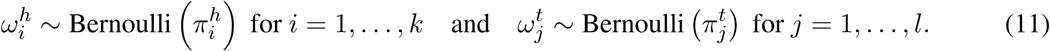
Where 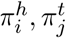 are posterior probabilities proportional to the respective alignment scores, as provided by the aligner. *Second*, 
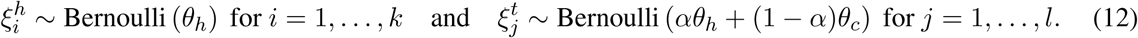

This reflects that sampling a fragment from the locus that is affected by the variant agrees with the probability to sample a genome copy affected by the variant from the bulk of cellular DNA. This, in turn, translates into the VAF *θ* of the variant in the respective sample^2^. We recall eq. (1) for the relationship between *θ_t_* and *θ_c_*. Whether *ξ_i_* is 1 or 0 is generally not evident from the observed *Z_i_*, due to typing uncertainty. We formally define 
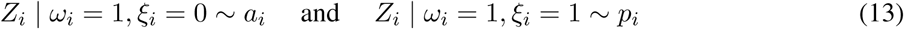
where *a_i_*(·) and *p_i_*(·) are the probability distributions of correct *Z_i_* when the indel is either *a*bsent or *p*resent. If *ω_i_* = 0, that is the alignment is incorrect, *Z_i_* | *ω_i_* = 0 ∼ 1 reflects that *Z_i_* has no influence on the posterior probability distribution of *θ*.

#### Observable Data: Split-Read and Internal Segment Alignments

We further specify distributions *a_i_* and *p_i_* for paired-end read alignments *Z_i_*. Note that, in general, our model does neither depend on paired-end alignments nor on a particular sequencing technology; see the Discussion for some final remarks on that point. Paired-end alignments can either overlap the variant locus with one of their read ends (*split-read alignments*) or with their unsequenced, internal segment (*internal segment alignments*).

For *internal segment alignments*, the *Z_i_* are integer-valued, reflecting the length of the alignment. Let *δ* denote the length of the indel at the locus in question, where *δ* is supposed to be negative for deletions and positive for insertions, and let *f*(·) specify the fragment length distribution for the sampled fragments. This distribution often turns out to be approximately Gaussian when following modern sequencing library protocols (e.g. [6]). Note however that our approach does not depend on the type of this distribution— any empirical fragment length distribution applies. When the alignment is aligned correctly and is not affected by the indel, *Z_i_* is governed by the fragment length distribution *f* itself. If the alignment aligns correctly and is affected by the variant, of length *δ*, the random variable *Z_i_* is governed by by *f_δ_*, defined by *f_δ_*(*z*) := *f* (*z* + *δ*). Note that in case of negative *δ* (= a deletion of length *δ*) this shifts *f* to the right, and vice versa for insertions. So, overall, 
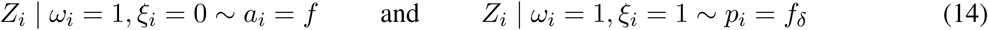
for internal segment alignments *Z_i_*. Since fragment length distributions are discrete, we evaluate *f* and *f_δ_* as 
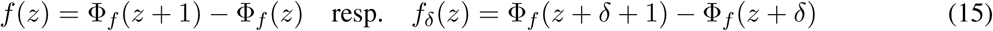
in our experiments, where Φ*_f_*(·) is the cumulative fragment length distribution function. Note that in our experiments, *f* is indeed approximately Gaussian, so we make use of the cumulative standard normal distribution function when evaluating Φ, where the necessary parameters *μ* and *σ* can be robustly estimated from the alignment data, see [14].

*Alignments Z_i_ whose ends overlap the variant locus* can either show a suitable split (i.e. a gap) or not. Further, depending on the read mapper, an indel can show up as a soft clip (i.e. a prefix or suffix of the read is not aligned to the reference). In order to become independent of the read mappers’s decisions, we strive to determine *a_i_* and *p_i_* by calculating the probabilty that the read has been sampled from either the reference or the alternative haplotype. For a given haplotype *H* and all reasonable offsets *O* of a read, we define this probability as 
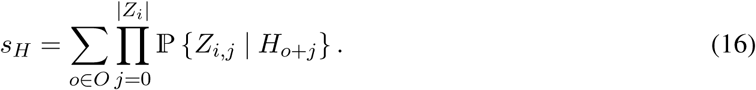

Here, unlike for segment alignments, *Z_i_* denotes the read sequence and 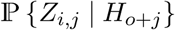 is the probability to observe base *Z_i,j_*· given that base *H_o_*_+_*_j_*·is the true allele (as defined by [3]). In other words, we calculate the probability that the read has been sampled from any position of the haplotype. In principle, an accurate calculation of *a_i_* and *p_i_* based on *s_H_* would require to consider all possible haplotypes implied by the combination of the variant and reference allele with possible surrounding variants within the range of the read (which can, e.g., be achieved via pair HMMs [4]). However, here we are only interested in the probability for the reference or alternative allele at the current locus, i.e., *s_r_* and *s_a_*. Since surrounding variants would impact both probabilities, they can be normalized away using the total probability, i.e., 
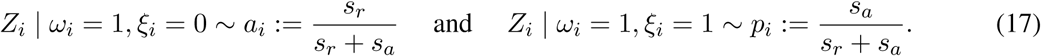

### Statements.

We finally obtain the Result outlined in the initial subsection 2.1 as a corollary to the following theorem.

### Theorem 2.2.

Let *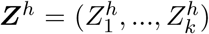, 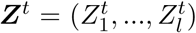* be the observable alignment data from a healthy and a tumor sample, covering the locus of a putative indel variant of length *δ*. Then 
- (i)The likelihood function 
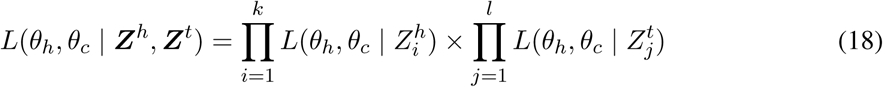
factors into likelihood functions referring to single alignments.
- (ii)Let *Z_i_* refer to any of the alignments, 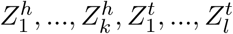 and let **ω*_i_*,**ξ*_i_* be its latent uncertainty hyperparameters. Then 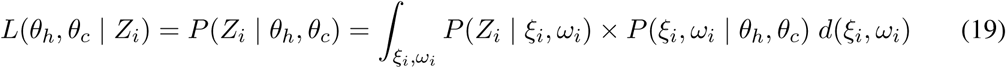

### PROOF

(*i*) follows immediately from the fact that the 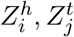 are conditionally independent given *θ_h_*,*θ_c_* see Figure 1b. (*ii*) follows from application of the Chapman-Kolmogorov equation, in combination with the dependency relationships captured by our model, see again Figure 1b.

### Corollary 1

*L*(*θ*_h_,*θ_c_*| ***Z**^h^, **Z**^t^*) *can be computed in O*(*k* + *l*) *runtime, with a small constant factor*.

The proof of Corollary 1 follows because the integral virtually is a sum over the three well-known cases {**ω*_i_* = 0}, {**ω*_i_* = 1, *ξ_i_* = 0} and {**ω*_i_* = 1, *ξ*_1_ = 1}, reflecting that the alignment *Z_i_* is either (1) incorrect, (2) correct and not affected by the variant, or (3) correct and affected by the variant, as previously mentioned. Because the proof is somewhat more technical, we have deferred it to subsection A in the Appendix.

## 3 Results and Discussion

### General Workflow

We present PROSIC (PROcessing Somatic Indel Calls) as a tool that implements the statistical model outlined in the Methods section. PROSIC requires a list of (putative) indel calls as VCF, and two BAM files, one of the cancer and one of the control genome. PROSIC then extracts somatic indel calls and estimates their VAF’s by implementing equations (5) [computing the probability of the putative indels to be somatic] and (10) [computing the maximum likelihood estimate (MLE) of the VAF’s] in subsection 2.1. See Figure 5, Appendix C for an illustration of the steps of PROSIC’s somatic indel calling pipeline: PROSIC implements 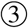 [computing probabilities to be somatic, eq. (5) and estimates of VAFs, eq. (10)] and 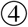 [FDR control], while 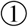 [read alignment] and 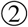 [generic indel calling] rely on existing state of the art, for which plenty of (often excellent) tools have been presented in the literature. Methods for 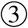 and 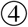, however, that transform the excellent generic indel callers into (equally excellent) somatic indel callers have been missing in so far. PROSIC is implemented in Rust [20], on top of the Rust-Bio library [10].

### Data

We used a real genome (Venter’s genome [13]), which has already been previously approved for NGS benchmarking purposes [17, 18] as a control genome and inserted randomly sampled 300,000 somatic point mutations, 150,000 insertions and 150,000 deletions, of which 279 509, 139 491 and 139 532 in the autosomes, respectively, following the clonal structure described by Figure 2 to obtain a simulated cancer genome. In terms of length, insertions and deletions follow the length distribution of Venter’s germline insertions and deletions. Reads were sampled using the Assemblathon read simulator SimSeq [5], at 30× and 40× for the control and the cancer genome, respectively. Subsequently, reads were aligned using BWA-MEM [14].

**Figure 2:**
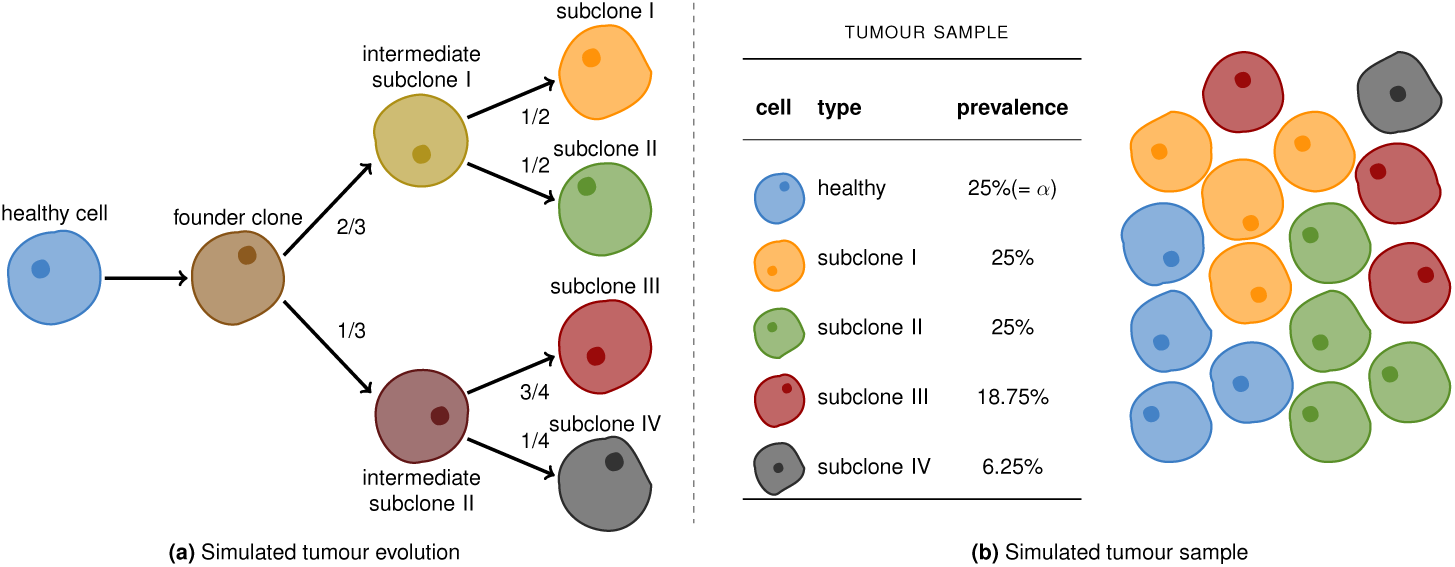
Simulated Cancer Clones: (**a**) The evolution of the cancer clones. (**b**) The simulated tumour sample. Each cell is assumed to be diploid. Cells of the same type share the same genetic code. The relative prevalences of the various cell types are shown in the table. The level of impurity (*α*) is 25%.

### Tools: Alignments and (Generic) Indel Callers

In our evaluation experiments, we exclusively focus on deletions. The reason is that none of the state-of-the-art (generic) indel calling tools yielded sufficient amounts of insertions of 30 bp and longer when applied to our simulated BAM files. Of course, once reliable insertion callers are available, PROSIC can be applied also there; as for insertions of 1-30 bp, PROSIC achieves performance rates roughly on a par with those achieved for deletions. For generating lists of indels in form of VCF files [step 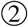 in Fig. 5, sec. C], we chose PINDEL 0.2.5 [30], PLATYPUS 0.8.1 [24] in assembly modus [since in default mode PLATYPUS does not discover indels longer than 30 bp], DELLY 0.7.6 [23] and LANCET 1.0.0 [22]. For PINDEL and PLATYPUS we applied an *ad-hoc routine* for discovery of somatic indels: we subtract indels called in the control genome from those called in the cancer genomes, and keep all those that pass auxiliary filters defined by the tools (PASS in the FILTER column). Both LANCET and DELLY provide an integrated ad-hoc method for *somatic indel discovery*. We provided the BWA-MEM alignments as input for all tools (which is often just the recommended choice of aligner, see LANCET, for example). When subsequently running PROSIC, we use the output VCF’s of the tools in combination with the BAM files that were the basis for generating the VCF’s.

### Experiments: Performance Rates

In the following, *Recall* is defined to be the fraction of true variants discovered, while *Precision* is the fraction of correctly predicted variants among the variants called overall. Figure 3 shows Recall and Precision for the tools from above on the simulated data, both in ad-hoc somatic variant calling mode and, in juxtaposition to this, when running PROSIC on their calls. PROSIC is consistently able to improve the recall of the corresponding ad-hoc calling approach, across all size ranges without impacting the precision. Moreover, because of the statistical framework, PROSIC allows to control the FDR, a highly favorable feature from a practical point of view, which cannot be warranted by ad-hoc routines.

**Figure 3:**
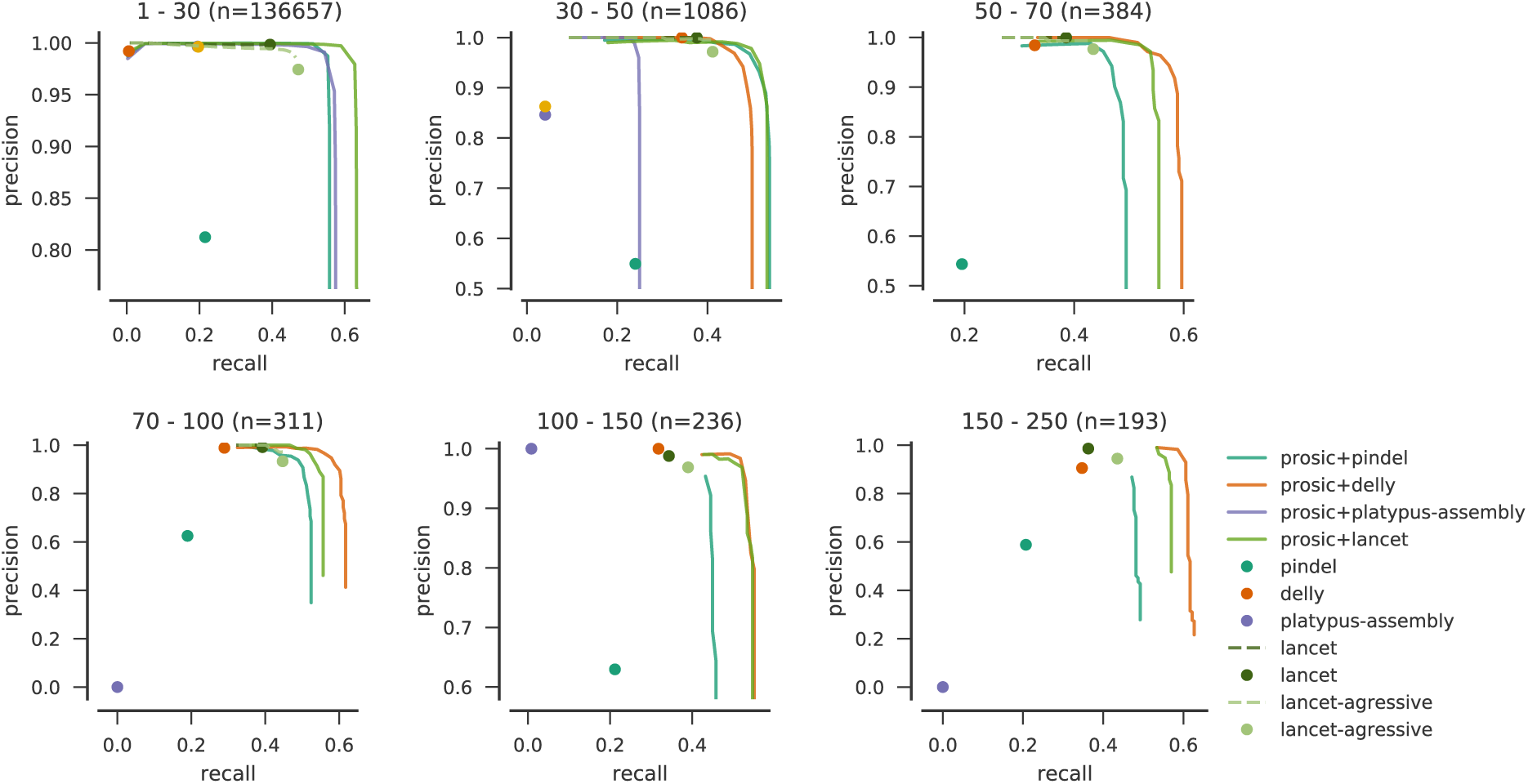
: Recall and precision for calling somatic deletions. For our approach (prosic+*) we controlled the FDR at increasing levels between 0.01 and 1.0 (resulting in a curve). For lancet, curves are plotted by scanning over the provided p-values (dashed curves). Ad-hoc results are shown as single dots. Note that PLATYPUS did not make considerable amounts of true predictions beyond 30bp, whereas DELLY did not provide considerable amounts of true positive calls smaller than 30bp.

### Experiments: Consistency of FDR Control

See further Figure 4 for consistency of FDR control levels in terms of precision achieved: it shows that our FDR control procedure indeed warrants the intended FDR, indicated by curves being above the dashed diagonal. In particular, FDR control is tight for reasonably small FDR values. For increasing FDR control levels, control becomes conservative.

**Figure 4:**
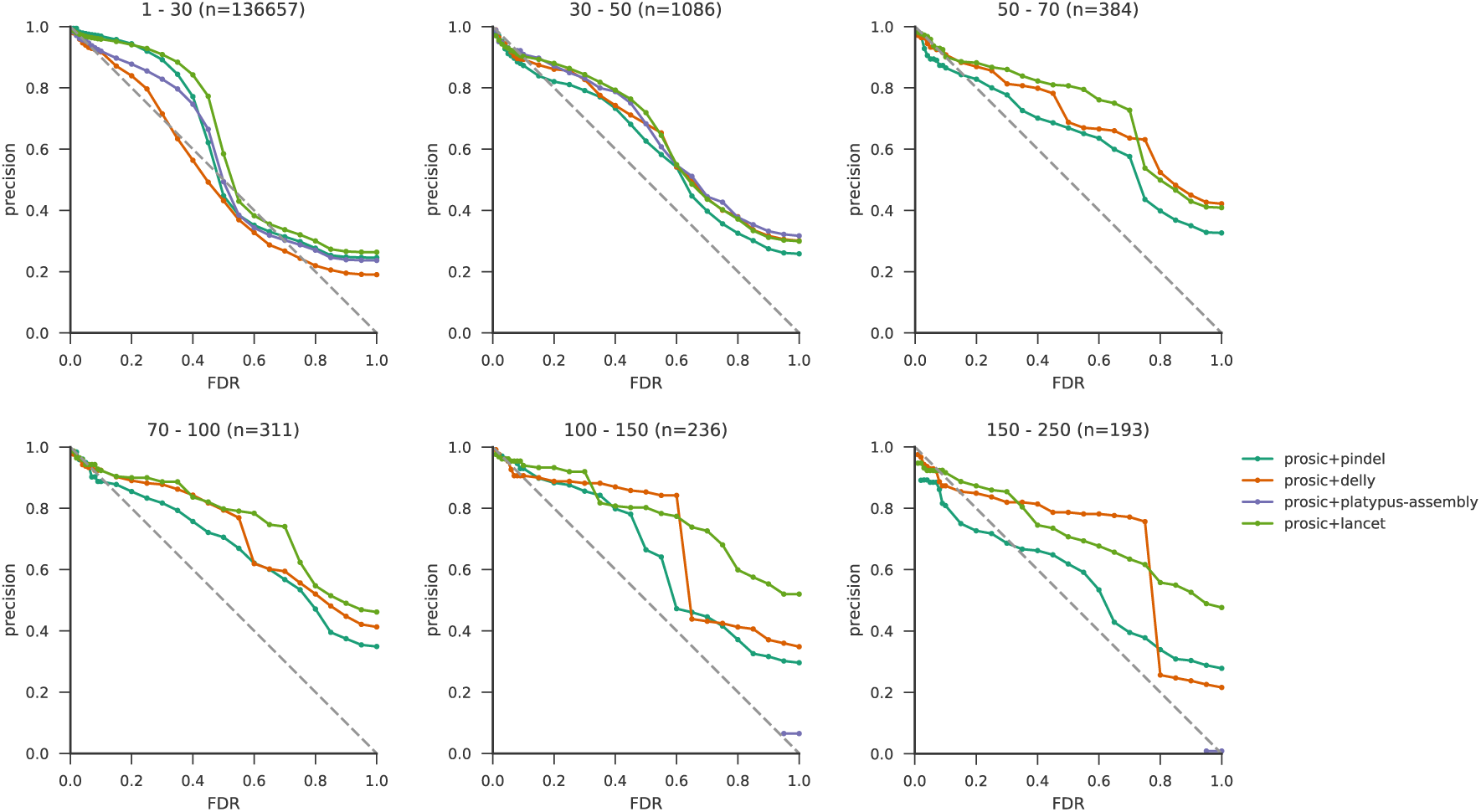
FDR control with PROSIC. For each FDR threshold, the corresponding precision is shown. A perfect FDR control would keep the precision curve exactly on the dashed diagonal. Above the diagonal, the control is conservative. Curves falling below the diagonal indicate underestimation of the FDR.

## Conclusion

We have provided a stastistical framework that allows to efficiently compute the likelihood of indel VAF’s given observed, yet (often heavily) uncertain alignment data from a cancer and a matched control genome. The efficiency in computation overcomes a principled computational bottleneck in uncertainty quantification and enables to compute 1) probabilities for indel variants to be somatic, 2) maximum likelihod estimates for their VAF’s and 3) reasonable, consistent FDR control levels. We have further shown that PROSIC, the corresponding tool, achieves substantial improvements over somatic indel calling routines offered by prevalent indel discovery tools. In addition to the improvements achieved, the FDR can be reliably controlled at all levels, which allows for utmost flexibility in somatic indel discovery experiments.

At last, note that our model also applies for third-generation sequencing (TGS) data—which come with their own uncertainty characteristics—as long as probabilities of the kind *P*(*Z_i_* | *ξ_i,ω_*),*P*(*ω_i_*) can be obtained from TGS aligners in constant time, through alignment scores and error profiles, which applies, at least for most prevalent classes of TGS data. In future work of ours, we are focusing on finetuning our model towards TGS data applications.

## A Proof of Corollary 1

We distinguish between alignments 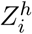 from the healthy sample and alignments 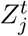 from the tumor sample. Making use of the above notation, we compute for 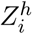, which does not depend on *θ_c_* (reflected by 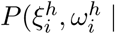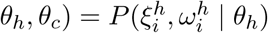 leading to the first equation) 
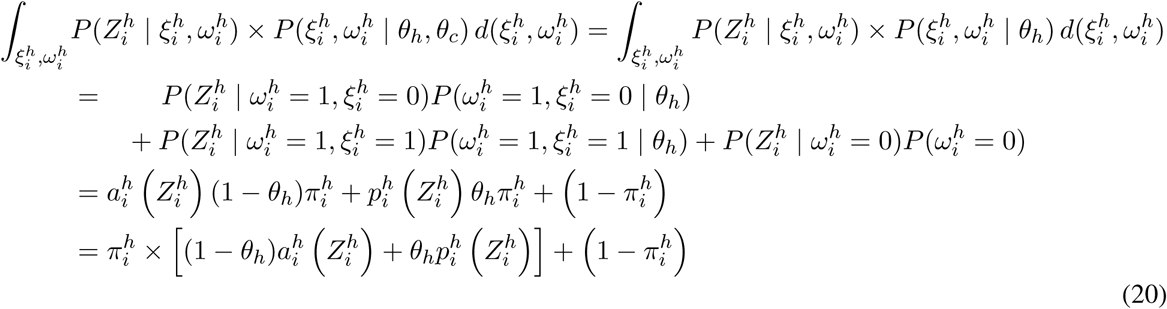
where in the last summand, the equation 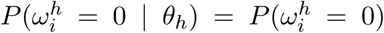 reflects that 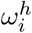 and *θ_h_* are independent, see Figure 1b. Analogously, while slightly more involved due to impurity considerations because 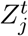 depends on both *θ_h_* and *θ_c_*, we compute

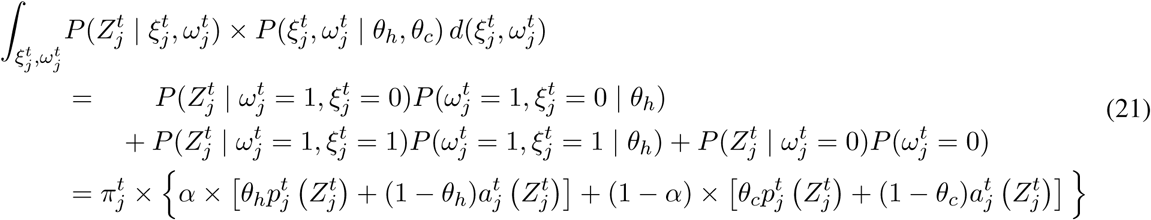

Note, at last, that all of the 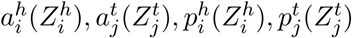 can be obtained in constant time and the amount of arithmetic operations required is small.

## B Uniqueness and computation of the maximum likelihood estimate

The likelihood function of *θ_h_* and *θc* given the data ***Z**^h^* and ***Z**^t^* as shown in eq. (2) is a higher-order polynomial, which makes it infeasible to derive its maximum analytically. We show in this section, however, that under weak conditions (as given in the following theorem) the likelihood function attains a unique global maximum on the unit interval for each value of *θ_h_*. We, in addition, show that the loglikelihood function is strictly concave, which simplifies the numerical maximization.

## Theorem B.1

The likelihood function L(θ_h_, θ_c_ | z^h^, z^t^) (where θ_h_ is fixed) attains a unique global maximum 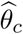 on the unit interval 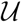 = [0, 1] when 
1. the likelihood of θ_h_ given the data from the healthy sample must be non-zero, i.e., 
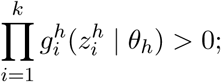
2. the subject 
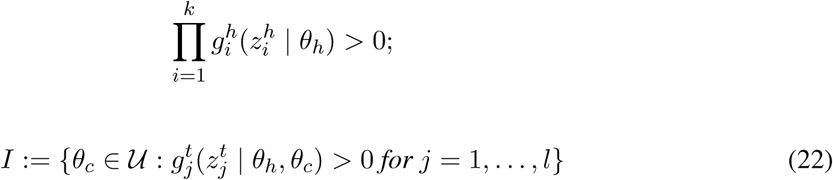
is connected and non-empty;
3. the level of impurity is smaller than 1, i.e., α < 1 (otherwise the ‘tumour’ sample would not contain any cancer cells);
4. there exists an observation 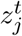 for which the alignment probability 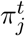 is strictly larger than zero and 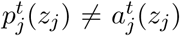 (i.e., there must exists an observation that with non-zero probability stems from the locus of interests and provided information about the presence or absence of the indel of interest).

*Proof*. The likelihood function with *θ_h_* fixed can be written in the form 
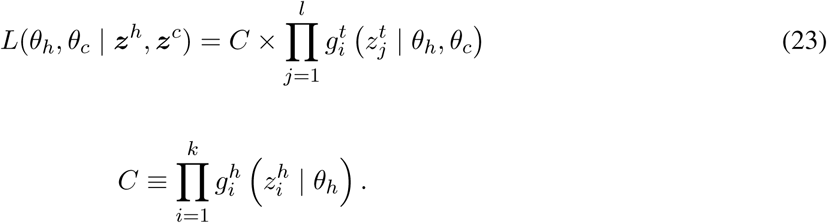
where *C* is the constant

In the case that condition (*1*) is *not* met, *C* = 0. The likelihood *L*(*θ_h_, θ_c_* | *z^h^, z^c^*) equals zero for all *θ_c_* and, therefore, does not attain a unique global maximum.

Suppose condition (*1*) is met (*C* > 0). Let us consider condition (*2*). Note that *L*(*θ_h_, θ_c_* | *z^h^, z^c^*) = 0 when *θ_c_* ∈ *I*, since for those *θ_c_* there exists an observation for which the likelihood 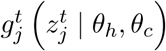 is equal to zero. The likelihood *L* is by definition strictly larger than zero when *θ_c_* ∈ *I*. Since the function 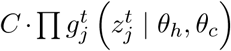 is a *l*-th order polynomial and, therefore, continuous, it must attain a global maximum on the interval *I*.

Suppose condition (*2*) is met. The point 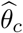 is a maximum of *L*(*θ_h_*, ·| *z^h^, z^c^*) iff it is a maximum of the loglikelihood function 
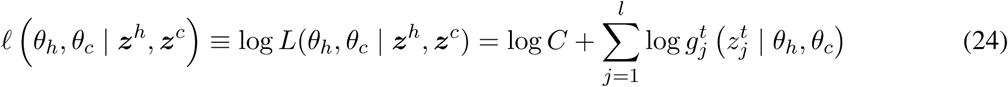
(with *θ_h_* fixed and *θ_c_* ∈ *I*) since the logarithm is a monotonic transform. (Note that *l* is only defined on the subset *I*). The second order derivative of the loglikelihood with respect to *θ_c_* is found to be 
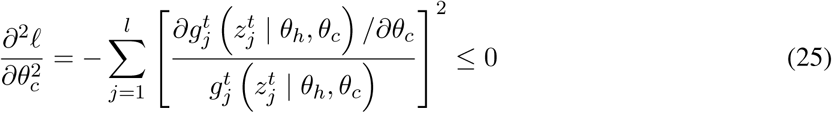
indicating that the loglikelihood function is concave. Note that it is strictly concave, i.e.,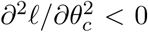, iff there exists an observation 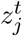 for which 
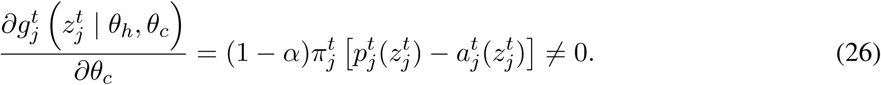

This inequality holds only when *α* ≠ 1, 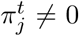 and 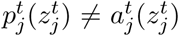, which constitutes conditions (*3*) and (4).

Suppose *I* is the non-empty closed set [*a, b*] on the unit interval. Since the loglikelihood is strictly concave when conditions (*3*) and (*4*) are met, it attains a unique global maximum 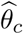 on *I*. Because the logarithm is a monotonic transformation, 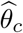 must be a unique global maximum of the likelihood function as well.

A similar reasoning holds when *I* is open or half-open. The maximum must lie on the interior of *I*, since the likelihood function is zero for those endpoints not in *I*. E.g., when *I* is the open interval (*a, b*), then *L*(*θ_h_*,*a* | *z^h^, z^c^*) = *L*(*θ_h_*,*b* | *z^h^, z^c^*) = 0 while *L*(*θ_h_*,*θ_c_* | *z^h^, z^c^*) is strictly positive on *I*. The loglikelihood function is under conditions (3) and (*4*) strictly concave on *I*, therefore, the likelihood function attains a unique global maximum.

We approximate the overall maximum of the likelihood function *L* by numerically maximizing (using Brent’s method) the likelihood function three times: and 1. We approximate the MLE of the VAFs by numerically maximizing the likelihood function *L*(*θ_h_, θ_c_* | *z^h^, z^c^*) using Brent’s method. where *θ_h_* takes the values 0, ½ and 1.

## C Workflow Figure

**Figure 5:**
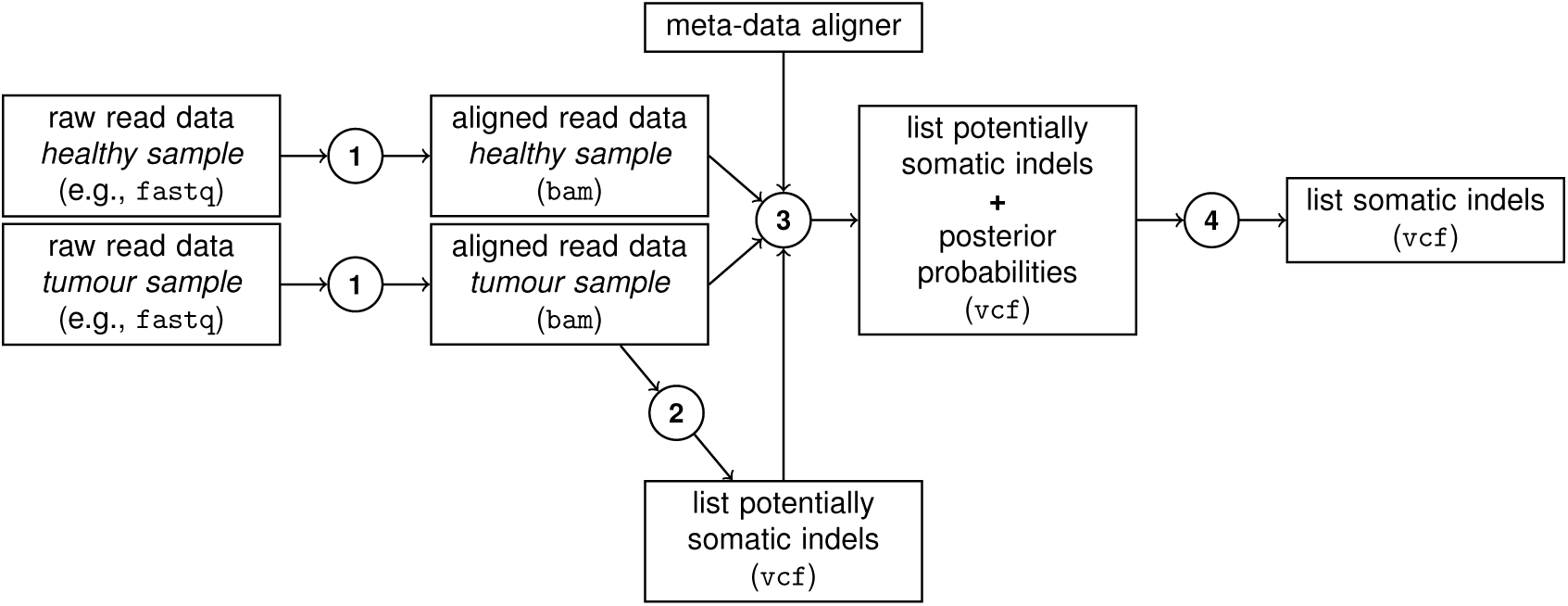
The overall pipeline for somatic indel calling presented in this paper. While (1) (= aligning short reads) and (2) (= calling indels) rely on existing (and plentiful available) state of the art, steps (3) [implementing eqs. (5), (10)] and (4) [FDR control] reflect methodology developed here.

## Footnotes

1 Here, we refer to insertions and deletions of all possible sizes, ranging from 1 to several thousands of base pairs.

2 Under the common assumption, of course, that chromosomes harboring and not harboring the indel are equally likely to bring forth a read.

